# Alteration in long noncoding RNAs in response to oxidative stress and ladostigil in SH-SY5Y cells

**DOI:** 10.1101/2022.02.20.481187

**Authors:** Keren Zohar, Eliran Giladi, Tsiona Eliyahu, Michal Linial

## Abstract

Microglia activation causes neuroinflammation, which is a hallmark of neurodegenerative disorders, brain injury, and aging. Ladostigil, a bifunctional reagent with antioxidant and anti-inflammatory properties, reduced microglial activation and enhanced brain functioning in elderly rats. In this study, we studied SH-SY5Y, a human neuroblastoma cell line, and tested viability in the presence of hydrogen peroxide and Sin1 (3-morpholinosydnonimine), which generates reactive oxygen and nitrogen species (ROS/RNS). Both stressors caused significant apoptosis and necrotic cell death that was attenuated by ladostigil. Our results from RNA-seq experiments show that long non-coding RNAs (lncRNAs) account for 30% of all transcripts in SH-SY5Y cells treated with Sin1 for 24 hours. Altogether, we identify 94 differently expressed lncRNAs in the presence of Sin1, including MALAT1, a highly expressed lncRNA with anti-inflammatory and anti-apoptotic functions. Additional activities of Sin-1 upregulated lncRNAs include redox homeostasis (e.g., MIAT, GABPB1-AS1), energy metabolism (HAND2-AS1), and neurodegeneration (e.g., MIAT, GABPB1-AS1, NEAT1). Four lncRNAs implicated as enhancers were significantly upregulated in cells exposed to Sin1 and ladostigil. Finally, we show that H_2_O_2_ and Sin1 increased the expression of DJ-1, a redox sensor and modulator of Nrf2 (nuclear factor erythroid 2– related factor 2). Nrf2 (NFE2L2 gene) is a major transcription factor regulating antioxidant genes. In the presence of ladostigil, DJ-1 expression is restored to its baseline. The mechanisms governing SH-SY5Y cell survival and homeostasis are highlighted by the beneficial role of ladostigil in the crosstalk involving Nrf2, antioxidant transcription factor DJ-1, and lncRNAs. Stress-dependent induction of lncRNAs represents an underappreciated regulatory level that contributes to cellular homeostasis and the capacity of SH-SY5Y to cope with oxidative stress.

## Introduction

Cellular homeostasis is a hallmark of maintaining normal brain function. A decline in cognitive function occurs along with normal brain aging. In humans and rodents, it includes the reduction of synaptic sites, enhanced apoptosis, alteration in the shape and number of dendritic spines, and extensive cell death [1,2]. The young brain successfully copes with stress and toxicity (e.g., oxidative stress), a capacity that drops in the aging brain [3,4]. Failure in the effectiveness of the stress response leads to pathological processes as documented in brain injury, ischemia and reperfusion (I/R), as well as neurodegenerative diseases [5,6].

The aged brain is signified by an elevation in genes associated with inflammation caused by microglial activation that contributes to age-dependent learning deficits [7,8]. In previous studies, old rats were chronically treated with ladostigil, a bifunctional reagent with antioxidant and anti-inflammatory activities [9,10]. The benefit of such treatment in restoring memory loss argues that ladostigil positively affects neurogenesis and cellular homeostasis [11,12]. In addition, paradigms based on acute insults for disturbing brain function (e.g., brain injury, hypoxia, viral infection) were developed, resulting in immediate changes in gene expression. At the molecular level, events that underlie proteotoxicity and mitochondrial malfunction were demonstrated by changes in the levels of gene expression, mostly of coding genes and miRNAs. A gradual change in gene expression levels also occurs throughout the progression of neurodegenerative diseases [13,14].

At the cellular level, the ability of cells to dissipate damage from external insults is carried out by the mitochondrial function and the antioxidants. Elevated levels of reactive oxygen and nitrogen species (ROS/RNS) damage cells through the modifications of DNA, lipids, and proteins [15]. Neurons that are exposed to damaging signals initiate a set of responses such as and alteration in ATP production, oxidative stress response by induction of antioxidants, apoptosis, and autophagy. The endoplasmic reticulum (ER) stress response allows cells to trigger autophagy for maintaining proteostasis [16,17]. All these mechanisms serve for dissipating the initial insult and controlling the damage [18,19].

In this study, we assayed the molecular events induced by extreme conditions (e.g., neurotoxicity or protein misfolding) by monitoring the gene expression profiles of SH-SY5Y cells, a neuroblastoma-derived cell line that carries some dopaminergic neuronal features [20]. We produced conditions that resemble abrupt (by H_2_O_2_) and continuous (by Sin1; 3-morpholinosydnonimine) oxidative stress conditions, the latter being a stressor that elicits prolonged oxidative stress. We then tested the effect of ladostigil on the molecular profile and viability of undifferentiated SH-SY5Y cells. Specifically, we focused on the role of lncRNAs under conditions of cell perturbations. In recent years, emerging evidence hints at the involvement of lncRNAs in neurodevelopmental, metabolic, and neurodegenerative diseases like Alzheimer’s disease (AD) [21,22]. However, the mode of action of lncRNA in cellular homeostasis remains unclear. Herein, we present an unbiased view on lncRNA and their induction following treatment with Sin1 and ladostigil. Our investigation of gene expression levels along with the attenuation of cell death by ladostigil suggests an overlooked role of lncRNAs in coping with cellular stress.

## 2. Materials and Methods

### 2.1. Materials

All reagents were purchased from Sigma Aldrich (USA) unless otherwise stated. Ladostigil, (6-(N-ethyl, N-methyl) carbamyloxy)-N propargyl-1(R)-aminoindan tartrate, was a gift from Spero BioPharma (Israel). Sin1 (3-(4-5 Morpholinyl) sydnone imine hydrochloride (Sigma-Aldrich, Cat-M5793). Media products MEM and F12 (ratio 1:1), heat-inactivated fetal calf serum (10% FCS), and L-Alanyl-L Glutamine were obtained from Biological Industries (Beit-Haemek, Israel). All tissue culture materials were purchased from Beit-Haemek (Israel). H_2_O_2_ (Sigma, Germany) was diluted in cold water and the solutions were used within 2 hrs. MTT (3-(4,5-dimethylthiazol-2-yl)-2,5-diphenyltetrazolium bromide) was used for cell viability (Sigma, Israel).

### 2.2. SH-SY5Y cell culture

Human neuroblastoma derived SH-SY5Y cells were obtained from ATCC (American Type Culture Collection, MD, USA) [23]. Cells were cultured in Minimum Essential Media (MEM and F12 ratio 1:1, 4.5 g/l glucose) with 10% fetal calf serum (FCS) and 1:10 L-Alanyl-L-Glutamine. Cells were incubated at 37°C in a humidified atmosphere of 5% CO_2_. Ladostigil was added to the culture medium 2 hours (hrs) prior to the activation of oxidative stress and analysis was performed Cells were tested 24 hrs later (i.e., 26 hrs after the addition of ladostigil), unless otherwise stated.

### 2.3. Cell viability assay

SH-SY5Y cells were cultured at a density of 2×10^4^ cells per well in 96-well plates, in 200 µl of medium. An MTT assay used colorimetric measurement for cell viability, with viable cells producing a dark blue formazan product. Cells were treated in the absence or presence of ladostigil concentrations (5.4 mM and 54 mM). The drug was solubilized in water. MTT solution in phosphate-buffered saline (PBS, pH 7.2) was prepared at a working stock of 5 mg/ml. After 24 hours, culture medium was supplemented with 10 µl of concentrated MTT per well. Absorption was determined in an ELISA-reader at *λ* = 535, using absorption at 635 nm as a baseline. Cell viability was expressed as a percentage of untreated cells. Each experimental condition was repeated 8 times. Results from survival assays are presented as mean ± SEM. When appropriate, p-values <0.05 were calculated and considered statistically significant. We report on genes that met FDR adjusted p-value of ≤0.05 (q-value).

### 2.4. Flow cytometry

SH-SY5Y cells were cultured in 6-well or 12-well plates to reach a 70-80% confluence level (37°C, 5% CO_2_) before analysis by fluorescence-activated single cell sorting (FACS). Increased fluorescence of Propidium iodide (PI) occurs upon binding to DNA. PI was used as a direct marker for permeable membranes of dead cells. PI and Annexin V-FITC were purchased from MBL (MEBCYTO-Apoptosis Kit). Conjugated Annexin V was used to detect early cell apoptosis by the increased intensity of the fluorescence signal. FACS analysis was done with 50,000 cells and the fraction of dead cell (PI), apoptotic cells were stained by Annexin V according to the manufacturer’s MBL protocol.

### 2.5. RNA sequencing

SH-SY5Y cells were pre-incubated for 2 hrs with ladostigil at a concentration of 5.4 μM. Cells were exposed to Sin1 (t=0) and harvested at 10 hrs or 24 hrs following the addition of ladostigil. Total RNA was extracted using the RNeasy Plus Universal Mini Kit (QIAGEN, GmbH, Hilden, Germany) according to the manufacturer’s protocol. Briefly, a mini spin column was used and centrifuged for 15s at ≥8000g at room temperature. Samples with RNA Integrity Number (RIN) > 9.0, as measured by Agilent 2100 Bioanalyzer, were considered for further analysis. Total RNA samples (1 μg RNA) were enriched for mRNAs by pull-down of poly(A) RNA and libraries were prepared using the KAPA Stranded mRNA-Seq Kit (QIAGEN, GmbH, Hilden, Germany) according to the manufacturer’s protocol. Since the RNA-seq experiment was performed using a protocol for the removal of ribosomal RNAs, restricting molecule length to a minimum of >200 nucleotides, this allowed to achieve sequencing depth of many of the lowly expressing transcripts. RNA-seq libraries were sequenced using Illumina NextSeq 500 to generate 85 bp single-end reads. RNA-seq data files were deposited in ArrayExpress [24] under accession number E-MTAB-10450.

### 2.6. Bioinformatic analysis and statistics

All next-generation sequencing (NGS) data underwent quality control using FastQC [25] and were processed using Trimmomatic [26]. All genomic loci were annotated using GENCODE version 37, aligned to GRCh38 using STAR aligner [27] using default parameters. All statistical tests were performed using R-base functions. Figures were generated using the R package ggplot2. The experiments contained a minimum of three biological replicates. Trimmed mean of M-values (TMM) was used for the normalization of RNA read counts. A TMM value of >4 was used as the lower cutoff value for gene expression levels. For quantifying relative expression, the sum of TMM values from all lnCRNA (>4) were considered 100% and only ncRNAs representing >0.5% of that total TMM are reported. Differential expression analysis was performed using edgeR [28]. Unless otherwise stated, genes with false discovery rate (FDR) adjusted p-value ≤0.05 and an absolute log fold-change (FC) above 0.3 were considered as significantly differentially expressed.

## 3. Results

### 3.1 Ladostigil suppresses necrotic cell death

Brain aging is accompanied by drastic changes in the cells’ transcriptomes of different brain regions [11]. The complexity of the aging brain and the ongoing changes in cell composition (e.g., the ratio between neurons and glia) limits the utility of bulk transcriptomic data for the tracing of cell-specific molecular alterations. To simplify the experimental system, we analyzed homogenous neuroblastoma-derived SH-SY5Y cells and tested their expression profile by RNA-seq under varying oxidative stress conditions [29].

**Figure 1** shows the results of flow cytometry to determine the fraction of necrotic cells (each experiment covered 50,000 cells). Propidium iodide (PI) was used as a marker for permeable membranes of dead cells. We observed a marked increase in necrosis, increasing from 5.4% in the absence of H_2_O_2_ (baseline) to 20.5% in the presence of a moderate H_2_O_2_ concentration of 80 μM. This increase in necrotic death occurred in cells exposed to two doses of H_2_O_2_ during the 24 hours of the experiment (at 3 and 18 hours). In the presence of ladostigil, the fraction of necrotic cells was reduced substantially to 11.4%, compared to 7.7% at baseline. We therefore conclude that ladostigil protects cells against non-reversible cell death.

**Figure 1.**
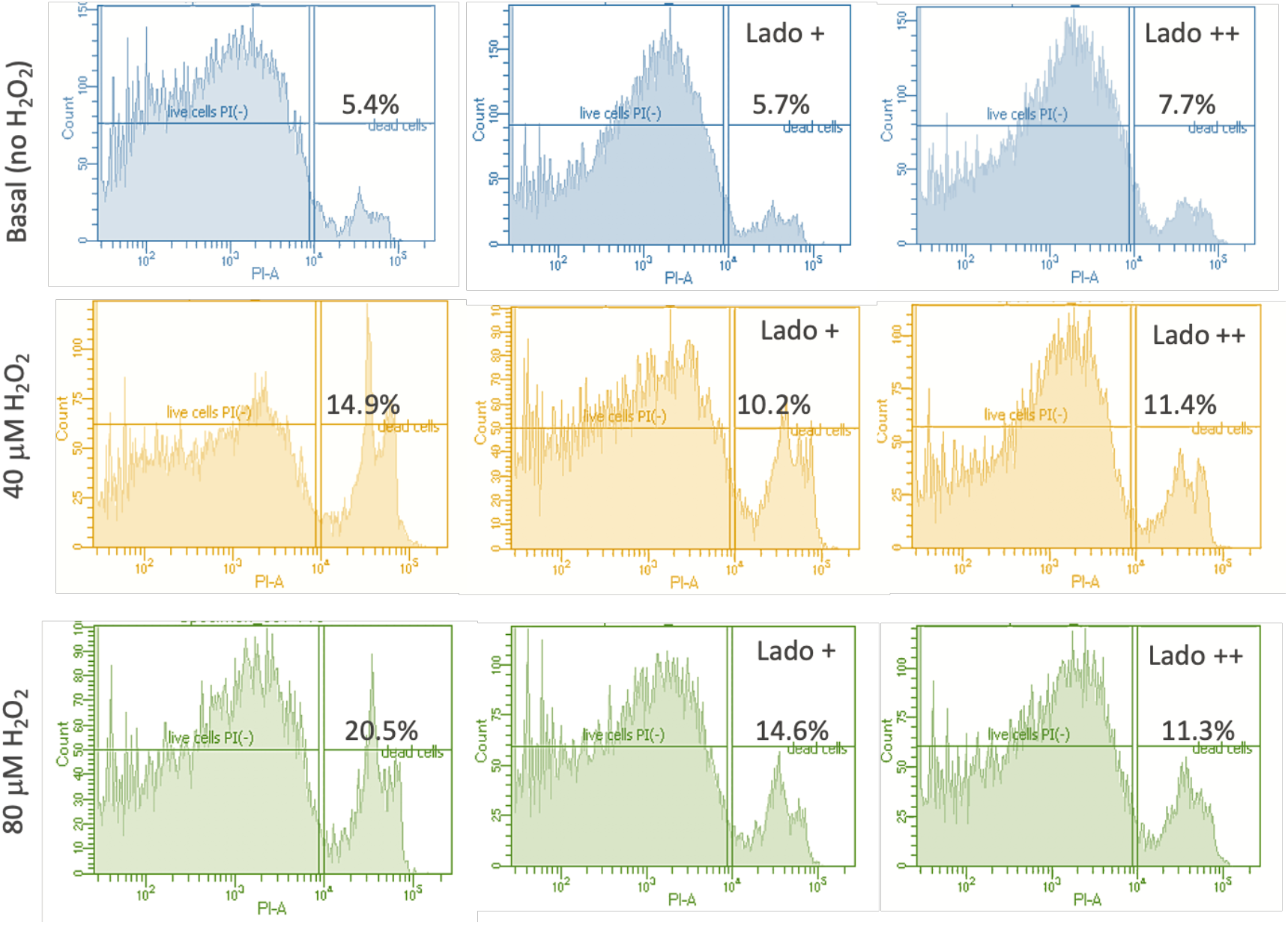
Flow cytometry analysis in naïve (untreated) and cells under oxidative stress condition in response to treatment with ladostigil (24 hrs). In each condition, 50k cells were tested and split to live and death cell in basal condition (blue), following 40 μM and 80 μM H2O2 (yellow and green panels, respectively), and by the addition of ladostigil at 5.4 μM (middle), 54 μM (right). The high fluorescence of PI (x-axis, above the vertical line) indicates the fraction of dead cells (in % from total).

### 3.2 Apoptotic signal in SH-SY5Y cells exposed to Sin1

To better mimic a prolonged and steady increase in oxidation, we exposed the cells to Sin1 at varying concentrations. Sin1 in living cells acts as a peroxynitrite donor (ONOO−) and generates nitric oxide (NO) and superoxide (O_2_−). The effect of prolonged oxidative stress produced by high levels of Sin1 on cell viability was assessed by flow cytometry using PI **(Figure 2A)**. Cell viability was tested at 5 hours following an increased concentration of Sin1. Under these harsh conditions, we also stained SH-SY5Y cells with Annexin-V, a sensitive early apoptosis marker. The fluorescence of Annexin-V detects the presence of phosphatidylserine (PS) at the outer leaflet of the plasma membrane. Staining cells with Annexin-V showed that the fraction of apoptotic cells is rather stable despite an elevation in overall cell death (**Figure 2B**). We propose that changes in the Sin1-dependent gene expression profile reflect the shift in the pathways driving cell death.

**Figure 2.**
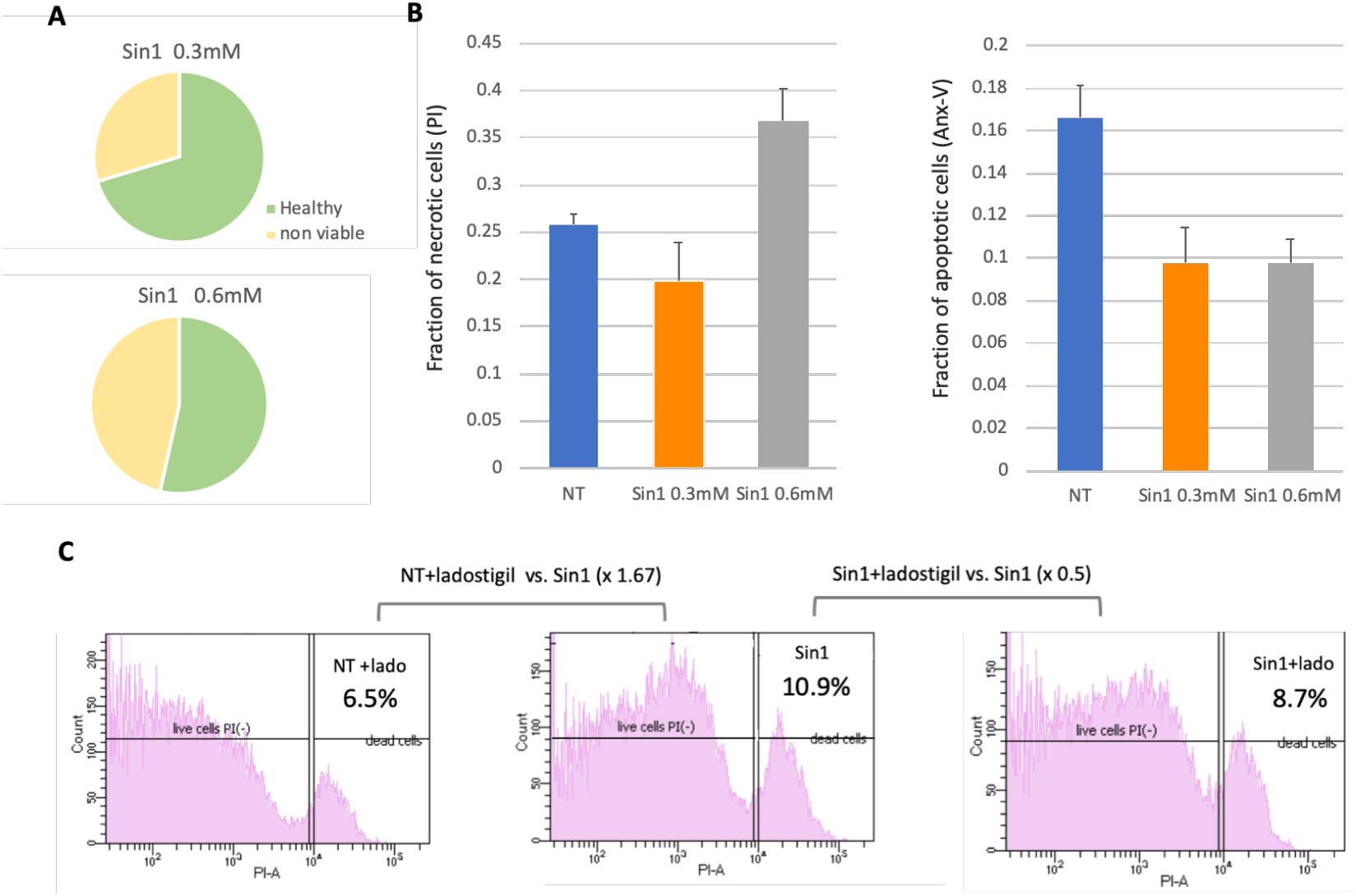
Apoptotic and necrotic cell death following oxidative stress. **(A)** Partition to healthy and non-viable cells from flow cytometry of SY-SY5Y cells treated with Sin1. **(B)** Results from 3 repeated experiments (mean and standard deviation) from flow cytometry run of SY-SY5Y cells treated with Sin1 for 5 hrs. The histogram summarizes the fraction of dead (left) and apoptotic (right) cells. **(C)** Flow cytometry analysis by of 50k cells following staining with PI. The high fluorescence of PI (x-axis, above the vertical line) indicates the fraction of dead cells (in % from total). Comparing of naïve (untreated, incubated with 5.4 μM ladostigil; left), upon oxidative stress induced by Sin1 (middle), and with Sin1 and treatment with ladostigil (right). As a control, we used incubated the cells with ladostigil without additional stressors (left) which had negligible effects on cell viability (24 hrs post incubation).

To test the potential of ladostigil (5.4 mM) to cope with prolonged mild stress by Sin1 (50 μM, 24 hours), we conducted flow cytometry analysis in cells without any induced stress (basal cell death, 6.5% of 50,000 living cells). The exposure to Sin1 increased the fraction of dead cells to 10.9%, a 1.7-fold increase. In the presence of ladostigil (5.4 mM), a 2-fold reduction in the fraction of dead cells relative to the Sin1 baseline **(Figure 2C)** was observed.

### 3.3. LncRNA expression levels in SH-SY5Y cells

In recent years, it has become evident that noncoding RNAs (ncRNAs) and specifically lncRNAs, play a role in cell homeostasis and the regulation of cell death. **Figure 3A** shows that cells’ transcriptome (RNA length >200 nt) comprises 70% mRNAs of coding genes (13,626 coding mRNAs out of a total of 19,475 transcripts), with the rest of the transcripts being partitioned among many ncRNA biotypes. The dominant class is lncRNA (long ncRNA, 63.2% of all ncRNAs), followed by pseudogenes (28.8%). However, the expression levels measured by TMM for most of these transcripts are very low (<4 TMM for biological triplicates, see Materials and Methods). By considering only transcripts that satisfy this threshold, we focused on 780 ncRNAs expressing transcripts that account for 6.7% of all transcripts (total of 11,588, **Figure 3B**). The rest of the analysis will consider only these ncRNAs (Supplementary **Table S1**). A subdivision according to biotypes confirms that most ncRNA transcripts belong to the lncRNA group (73%), followed by the unprocessed pseudogenes group (19%, **Figure 3B). Figure 3C** shows the top-expressing ncRNAs in SH-SY5Y cells (TMM >100). The division of these highly expressed transcripts into functional classes reveals an enrichment of mitochondrial transcripts, with MT-RNR1 and MT-RNR2 dominating (12S and 16S rRNAs of the mitochondrial ribosome, respectively). The importance of mitochondrial activity in SH-SY5Y cell viability is verified by the high expression level of mitochondrial enzymes (e.g., cytochrome c oxidase, NADH ubiquinone oxidoreductase; Supplementary **Table S1**). While the expression levels of MT-RNR1 and MT-RNR2 are exceptionally high, MT-RNR2 expression is 13-fold relative to MT-RNR1 expression. It was proposed that MT-RNR2 in addition to its role in ribosome formation is a source for short active peptides that have been linked to neuroprotection and Alzheimer’s disease (AD) [30].

**Figure 3.**
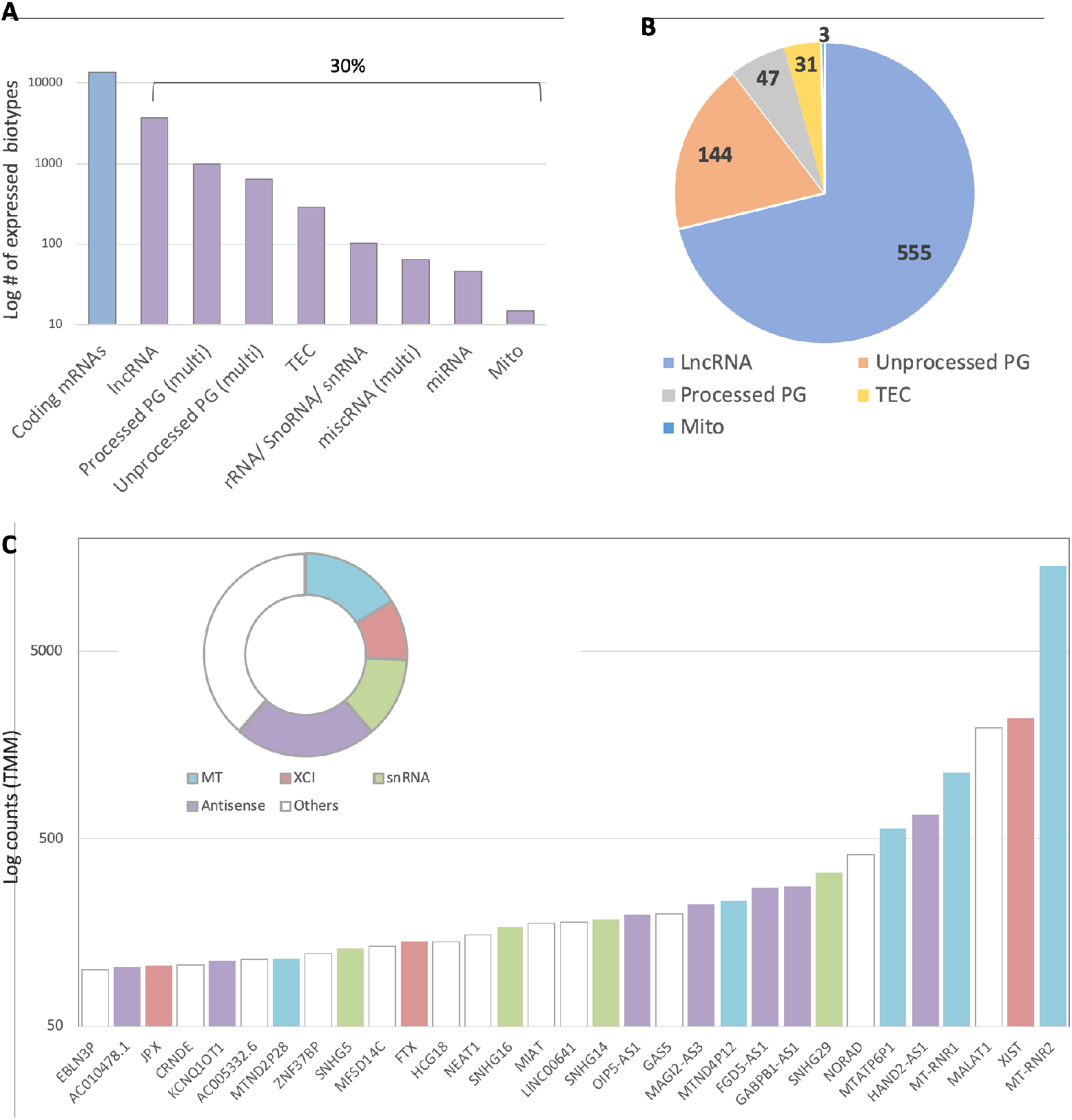
Partition of ncRNA biotypes in SH-SY5Y cells. **(A)** Partition of the RNA-seq results of untreated SH-SY5Y cells according to the major molecular biotypes. Altogether there are 19,475 identified transcripts. Blue, coding genes; purple, ncRNAs portioned by biotype classes. Mito, mitochondrial transcript; TEC, to be experimentally confirmed; PG, pseudogene. Combined related biotypes (e.g., transcribed unprocessed and unitary pseudogene) are marked as multi. **(B)** Partition of 780 expressed ncRNA transcripts partitioned by major classes. The ncRNA are based on transcripts with average TMM >3. There are 11,588 aligned transcripts that meet this threshold. **(C)** Top expressing ncRNA (TMM >100) according to their broad-sense annotation class and their partition (inset). MT, mitochondria; snRNA, small nucleus RNA; XCI, X-chromosome inactivation.

In addition, XIST, FTX, and JPX are ncRNAs that govern the X-chromosomal inhibition process in the female genome, supporting the role of abundant ncRNAs in chromatin compaction. In particular, many lncRNAs we found here (**Figure 3C**) have been implicated in the regulation of oxidative stress, particularly in the context of the CNS and neurons [31]. The listed highly expressed ncRNAs (denoted “others”) are less defined but share regulatory functions. Among these uncharacterized ncRNAs is MALAT1 (Metastasis-associated lung adenocarcinoma transcript 1; expression level ranked 99.94% of all ncRNA-expressed genes). The MALAT1 gene was shown to have a neuroprotective role in AD and Parkinson’s disease (PD) [32,33]. MALAT1 also protects hippocampal neurons against autophagy, apoptosis [34] and ischemia [35]. Other genes (**Figure 3C**) include NORAD, GAS5, LINC000641, MIAT, and NEAT1 (ranked from 97.49% to 88.37% of all expressed ncRNA genes). Downregulation of these lncRNAs was shown to prevent apoptotic cell death in the context of AD and PD [36]. NEAT1 and SNHG1 were implicated in stress-induced autophagy in PD [37] whereas MIAT promotes neural cell autophagy and apoptosis in ischemic settings [38]. Thus, the level of expression of highly abundant ncRNAs allows cells to alter apoptosis, autophagy, and necrotic cell death and cope with oxidative-induced damage.

### 3.4. Chronic oxidative stress upregulated a collection of lncRNAs

Ample studies have shown a correlation between oxidative stress and lncRNA expression [39]. Based on the results showing reduced cell viability by Sin1 and a beneficial effect of ladostigil (**Figure 2C**), we tested the profile of the cells’ ncRNA levels 24 hours following the Sin1 stimulus. **Figure 4A** shows the number of differentially expressed (DE) ncRNA transcripts partitioned to lncRNAs and other ncRNA types according to their expression profile. We found that the strongest effect induced by Sin1 is the upregulation of ncRNAs. The fraction of ncRNAs among DE downregulated genes is only 5.6%, while among the upregulation transcripts it accounts for 17.1% (p-value 1.2e-02). Moreover, among all upregulated ncRNAs, the lncRNAs class is strongly enriched. Among the downregulated genes, the ratio of lncRNAs to all other molecular ncRNA biotypes is 2.1 and 5.1 for the upregulated genes (**Figure 4A**).

**Figure 4.**
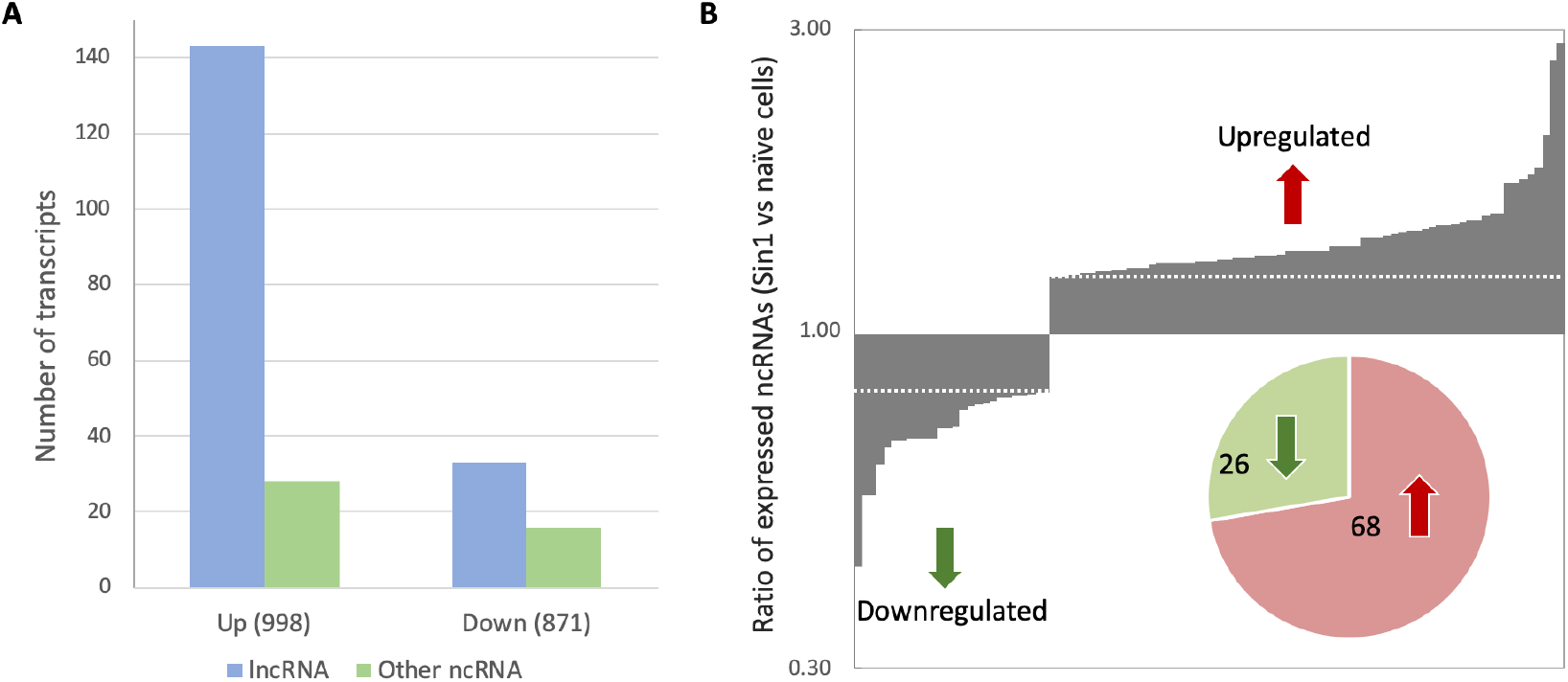
Differentially expressed (DE) ncRNAs following 24 hours exposure to Sin1 relative to untreated cells. **(A)** Transcripts associated with upregulation (Up) and downregulation (Down) relative to naïve, untreated cells. The number of genes associated with each expression trend is shown in parenthesis. The rest of the genes (9,719 genes) are unchanged. See Materials and Methods for expression trend definition. **(B)** Histogram showing the DE ncRNAs ratio for downregulated and upregulated transcripts. The dashed line marks the expression ratio (1.25 and 0.8) vs. naïve cells for Up and Down, respectively. The pie diagram applies to 94 genes that satisfy the thresholds (**Supplementary Table S3)**.

We conclude that the fraction of the expressed ncRNAs, and specifically the lncRNA class, significantly increased under Sin1-induced stress. While many coding genes are induced by Sin1 [29], the fold change in ncRNA expression was modest (bounded by <3-fold, **Figure 4B**). **Table 1** shows the list of lncRNAs following Sin1 treatment with a predetermined expression level (>0.5%, see Materials and Methods). All 18 Sin1 upregulated ncRNAs (**Table 1)** are statistically significant (p-value FDR <1.0e-3; Supplementary **Table S2**). The highest expressed lncRNA is MALAT1 (**Figure 3C**), which is also upregulated by Sin1 (**Table 1**, p-value FDR <2.5e-04).

**Table 1.** Differentially expressed lncRNA genes in cells exposed to Sin1 (24 hours)

| lncRNA | lncRNA – Full name | FDR | DE (fold) | TMM<sup>a</sup> |
| --- | --- | --- | --- | --- |
| MALAT1 | metastasis associated lung adenocarcinoma transcript 1 | 2.5e-04 | 1.24 | 1963.7 |
| HAND2-AS1 | HAND2 antisense RNA 1 | 6.7e-25 | 1.33 | 671.8 |
| GABPB1-AS1 | GABPB1 antisense RNA 1 | 1.7e-18 | 1.38 | 280.8 |
| MIAT | myocardial infarction associated transcript | 6.3e-21 | 1.35 | 176.9 |
| NEAT1 | nuclear paraspeckle assembly transcript 1 | 3.5e-05 | 1.49 | 154.0 |
| CASC15 | cancer susceptibility 15 | 1.5e-16 | 1.27 | 85.8 |
| AC093849.2 | novel transcript | 2.9e-33 | 1.46 | 85.0 |
| CCDC144NL-AS1 | CCDC144NL antisense RNA 1 | 1.6e-12 | 1.32 | 67.5 |
| MIR137HG | MIR137 host gene | 4.5e-23 | 1.47 | 58.7 |
| GABPB1-IT1 | GABPB1 intronic transcript | 4.6e-15 | 1.35 | 45.6 |
| LINC02268 | long intergenic non-protein coding RNA 2268 | 1.2e-16 | 1.45 | 44.6 |
| TTN-AS1 | TTN antisense RNA 1 | 1.2e-06 | 1.26 | 43.0 |
| LINC01833 | long intergenic non-protein coding RNA 1833 | 6.8e-14 | 1.32 | 42.0 |
| LINC00632 | long intergenic non-protein coding RNA 632 | 5.0e-08 | 1.31 | 40.2 |
| AL392083.1 | novel transcript, sense intronic to ADARB2 | 1.8e-33 | 1.78 | 37.6 |
| GATA3-AS1 | GATA3 antisense RNA 1 | 2.3e-08 | 1.32 | 36.5 |
| AC012354.1 | novel transcript | 2.1e-08 | 1.28 | 32.7 |
| AC015813.1 | novel transcript | 4.3e-07 | 1.35 | 31.8 |
| AC136621.1 | novel transcript (TEC) | 2.6e-27 | 0.56 | 37.7 |
| AL645608.2 | novel transcript | 4.7e-08 | 0.71 | 35.9 |
| AC010931.1 | novel transcript | 1.5e-19 | 0.68 | 34.9 |
| LINC01128 | long intergenic non-protein coding RNA 1128 | 1.2e-16 | 0.71 | 32.2 |
| EP400P1 | EP400 pseudogene 1 | 4.3e-06 | 0.78 | 32.1 |

<sup>a</sup>TMM, trimmed mean of M-values

Among the Sin1 induced lncRNAs, NEAT1 was upregulated by 1.49 fold (FDR p-value 3.5e-05). NEAT1 is a neuronal lncRNA that could reverse the damage caused by superoxide and counteract the H_2_O_2_-induced neuronal damage. Many of the listed lncRNAs transcripts may act as a sponge for miRNAs (‘miRNA sponging’), exemplifying the competitive endogenous RNA (ceRNA) hypothesis. In addition, some of the ncRNAs are considered direct and indirect regulators of Nrf2 antioxidant signaling and as targets that alters oxidative stress through miRNAs [40,41]. The high expression level of HAND2-AS1 was postulated in driving an imbalance in energy metabolism [42]. We conclude that Sin1 exposure induces a broad range of stress-related transcriptional regulations mediated by lncRNAs.

Notably, among the Sin1 upregulated genes many are annotated as antisense (21% of all transcripts upregulated by Sin1**; Supplementary Table S3)**. For example, the lncRNA GABPB1-AS1 suppresses the transcription of GABPB1, a gene product that acts upon damaged membranes via peroxidases. As such, it is a master regulator of the cellular antioxidant capacity and cell viability in health and disease [43].

**Table 1** also lists lncRNAs that are downregulated by Sin1 (5 transcripts, Supplementary **Table S3**). Note that the absolute expression levels of these downregulated genes are moderate. While AC136621.1 and AC010931.1 were mostly studied in the context of cancer tissues, LINC01128 was predicted to bind Argonaut (AGO) proteins and synaptojanin 1 (SYNJ1). The latter is a principal protein in maintaining membrane dynamics, cellular trafficking, and autophagy.

### 3.5. Cancer-related SNHG family members are differentially expressed by Sin1

Small nucleolar RNA host genes (SNHGs) comprise a large family of lncRNAs whose expression greatly fluctuates [44]. Many of these lncRNAs are upregulated across cancer types and implicated as oncogenes. The function of the studied SNHGs in cancer is ceRNA sponging of miRNAs and direct interaction with mRNA and proteins [45]. We observed 17 significantly expressed transcripts that account for 3.3% of total, significantly expressed lncRNAs (Supplementary **Table S4**). **Figure 5A** presents the expression levels (log_10_TMM) of the identified SNHGs and the fold change in expression levels following Sin1 treatment. SNHG3, SNHG12 and SNHG15 were significantly downregulated by Sin1. Of these, SNHG12, downregulated 1.5-fold in the presence of Sin1, participates in the unfolded protein response (UPR) in apoptosis, inflammation, and oxidative stress in SH-SY5Y cells [46]. It is proposed that several of the SNHG members carry anti-inflammatory roles under oxygen-glucose deprivation/reoxygenation (OGD/R) conditions by targeting specific miRNAs [31,47]. In such cases, the ratio of free cellular miRNAs can be slightly changed, leading to a revised cell state by the available miRNA pool [48]. Our observations emphasize the importance of SNHGs beyond cancer in cell physiology and homeostasis (**Figure 5**).

**Figure 5.**
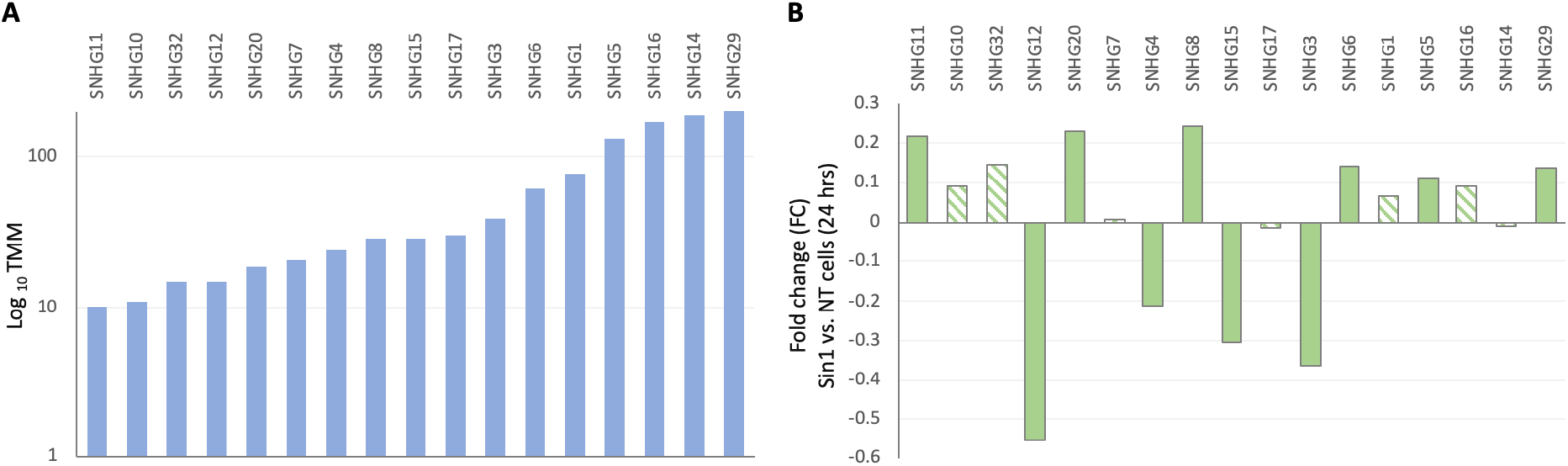
Expression of members of the small nucleolar RNA host genes (SNHGs). **(A)** The mean expression (by log10TMM) of the 17 SNHG members identified by DE lncRNA analysis. **(B)** The fold change (FC) following treatment with 50 μM Sin1 is shown for each of the SNHGs. The FC below and above zero shows downregulated and upregulation of the indicated genes. The striped color indicates genes belonging to the SNHG group but failed to reach FDR (p-value <0.05).

### 3.6. The impact of ladostigil on ncRNA profiles

**Figure 6A** shows four of the most significantly upregulated lncRNAs by Sin1 (p-value FDR ranges from 1e-23 to 1e-51) but unchanged in the presence of ladostigil (24 hrs). As a control, we incubated cells with ladostigil for 2 hrs which had negligible effects on cell viability (24 hrs post incubation, Supplementary **Table S1**). Remarkably, while only 8 genes (of 19,475 annotated genes) are significantly induced by ladostigil, four of them belong to lncRNAs **(Figure 6B**).

**Figure 6.**
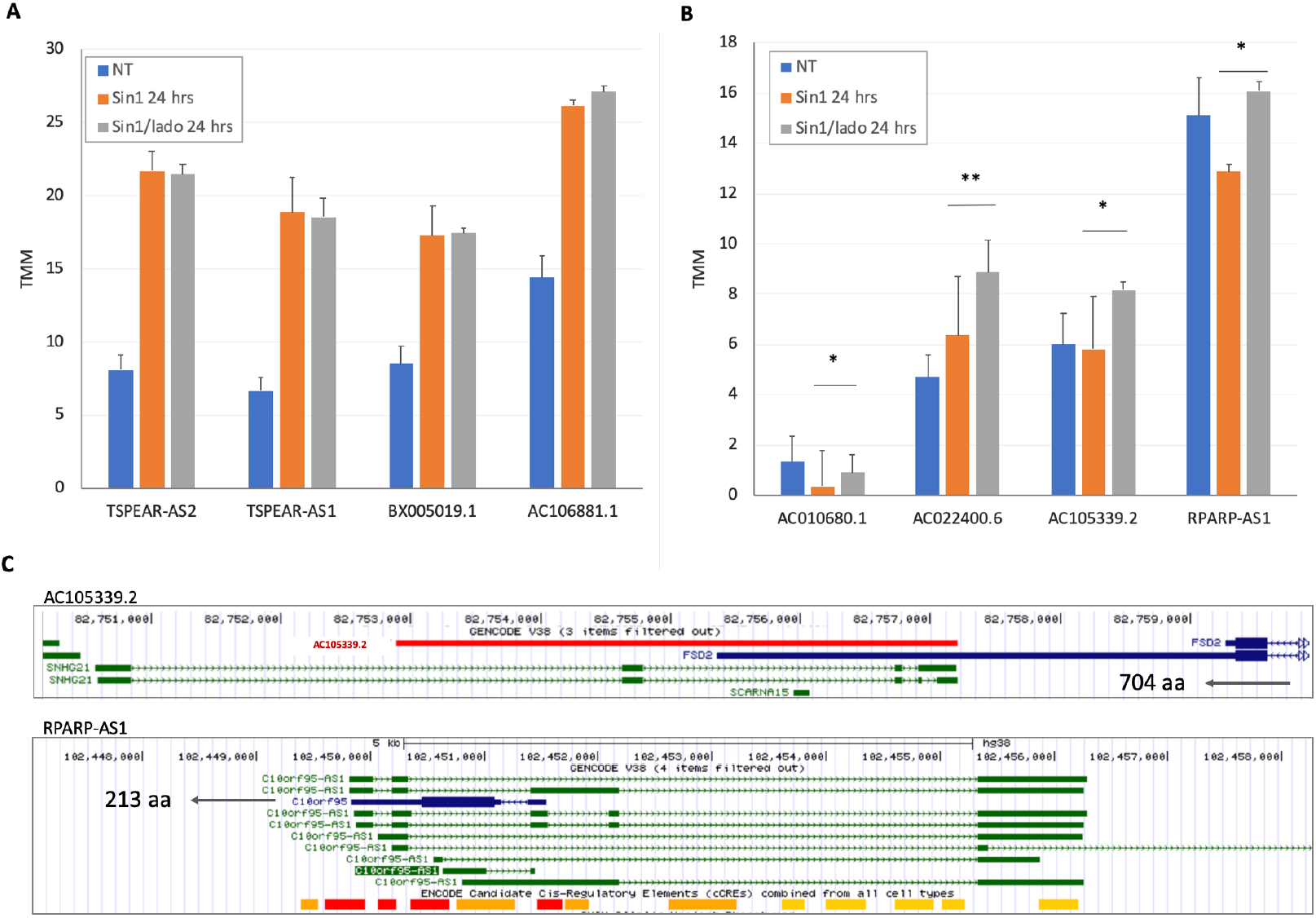
Sample of the lncRNAs by the measured expression levels in naïve untreated cells (NT), cells exposed to Sin1 (24 hrs) with or without ladostigil. **(A)** The transcripts shown are those with maximal induction by Sin 1 relative to NT. The average TMM and the standard deviations from biological triplicates are shown for each experimental group. **(B)** The transcripts shown are those with maximal effect by ladostigil in the presence of Sin1. The average TMM and the standard deviations from biological triplicates are shown for each experimental group. The statistical significance of p-value FDR ranging from 0.05-0.005 is depicted by * and <0.005 by **. **(C)** The genomic segment of AC105339.2 and RPARP-AS1 covers 10 and 12k nucleotides, respectively. AC105339.2 and RPARP-AS1 show the overlap between the lncRNA transcript and the gene in the vicinity. AC105339.2 acts as antisense to FSD2 (704 aa) and RPARP-AS1 is the antisense for C10orf95 (213 aa). Numerous transcripts of RPARP-AS1 are shown, above the ENCODE annotations for enhancers (red) and proximal promotors (yellow).

The RPARP-AS1 (also called C10orf95 Antisense RNA 1) overlaps with the production of a short (213 aa) protein of unknown function (**Figure 6C**). Possible regulation of the splicing machinery is proposed for AC105339.2 which overlaps with FSD2 transcript (fibronectin type III and SPRY domain containing 2; also called A1L4K1). The transcript that is induced by ladostigil most likely competes with the production of functional SFD2, which is involved in various RNA processing pathways. Two other overlapping genes, SNHG21 and SCARNA15 (**Figure 6C**) were studied in cancer tissues. For example, SCARNA15 plays a role in redox homeostasis and cancer cell survival [49].

We further inspected the genomic organization of these 4 ncRNAs upregulated by ladostigil in cells exposed to Sin1 (24 hrs). We used GeneHancer (GH) [50] to test the likelihood of these ladostigil upregulated ncRNAs to act as enhancers (eRNAs). GeneHancer integrates numerous five independent resources for enhancers. Elite enhancers are defined by high confidence scores.

**Table 2** presents the prediction confidence for the four ladostigil-upregulated ncRNAs as enhancers in view of the most likely regulated gene targets. Interestingly, while only 7% of the >100k analyzed genes and transcripts in the GH database are marked as elite enhancers, all genes found in our study are annotated as such. Two of the four transcripts were labeled as GH double elite. We further analyzed the potential of these double elite ncRNAs given the cellular response to ladostigil. The AC022400.6 transcript acts as an enhancer for 13 ncRNAs and 10 coding genes. Among these genes is SEC24C, which was implicated in the regulation of ER stress and mitochondrial redox [51]. SEC24 is part of ER stress induced autophagy. However, it was mostly studied in yeast. In the brain, SEC24C was previously validated as an essential protein for neuronal homeostasis by controlling the proper sorting of secretory vesicles [52]. The other double elite GH enhancer is RPARP-AS1, which targets 18 genes, including peroxisome proliferator-activated receptor gamma, coactivator-related 1 (PPRC1). This coactivator augments mitochondrial respiratory capacity and is involved in the transcription coactivation of Nrf1 target genes. Under stress, ladogtigil induction of lncRNAs may impact processes such as membranous trafficking, and respiratory-associated transcription.

**Table 2.** ncRNA upregulated by ladostigil treatment under oxidative stress condition

| ncRNA | GH score | Total score | Elite GH<sup>a</sup> | TSS (kb) | DB evidence | TFs | Coding targets | ncRNA targets | Selected target |
| --- | --- | --- | --- | --- | --- | --- | --- | --- | --- |
| AC010680.1 | 0.3 | **250.7** | * | 64.0 | 1/5 | 1.2 | 2 | 3 | FKBP7 |
| AC022400.6 | **1.9** | **250.7** | ** | 486.4 | 5/5 | 4.6 | 10 | 13 | SEC24C |
| AC105339.2 | **2.1** | 11.3 | * | 24.1 | 5/5 | 9.8 | 7 | 5 | WHAMM |
| RPARP-AS1 | **2.1** | **257.3** | ** | 546.8 | 5/5 | 6.6 | 13 | 5 | PPRC1 |

<sup>a</sup>GeneHancer (GH) definition of elite (\*) gene based on enhancer score and/or high confidence for the gene targets. High confidence scores are shown in bold.

### 3.7. Nrf2 signaling under stress condition is attenuating by ladostigil

Nrf2 (gene name NFE2L2) is a master regulator of expression of antioxidative enzymes in response to oxidative stress [53]. Consequently, it can control the redox cell state and suppress neurotoxicity and apoptosis [54]. We tested DJ-1 (also called PARK7), a major upstream regulator of NRf2, and its responsiveness to oxidation stress in SH-SY5Y cells. Specifically, we quantified the changes in expression of DJ-1 under H_2_O_2_ or Sin1 induced stress, and in response to ladostigil. **Figure 7A** shows the amplicon of DJ-1 relative to β-actin as a control upon activation with H_2_O_2_ or Sin1, and its expression level in the presence of ladostigil. Both conditions of induced oxidative stress led to upregulation of DJ-1. However, in the presence of a high level of ladostigil, DJ-1 expression level was suppressed and did not surpassbaseline levels. The results were further substantiated by using qRT-PCR (**Figure 7B)** that quantified the relative expression of DJ-1 relative to β-actin.

**Figure 7.**
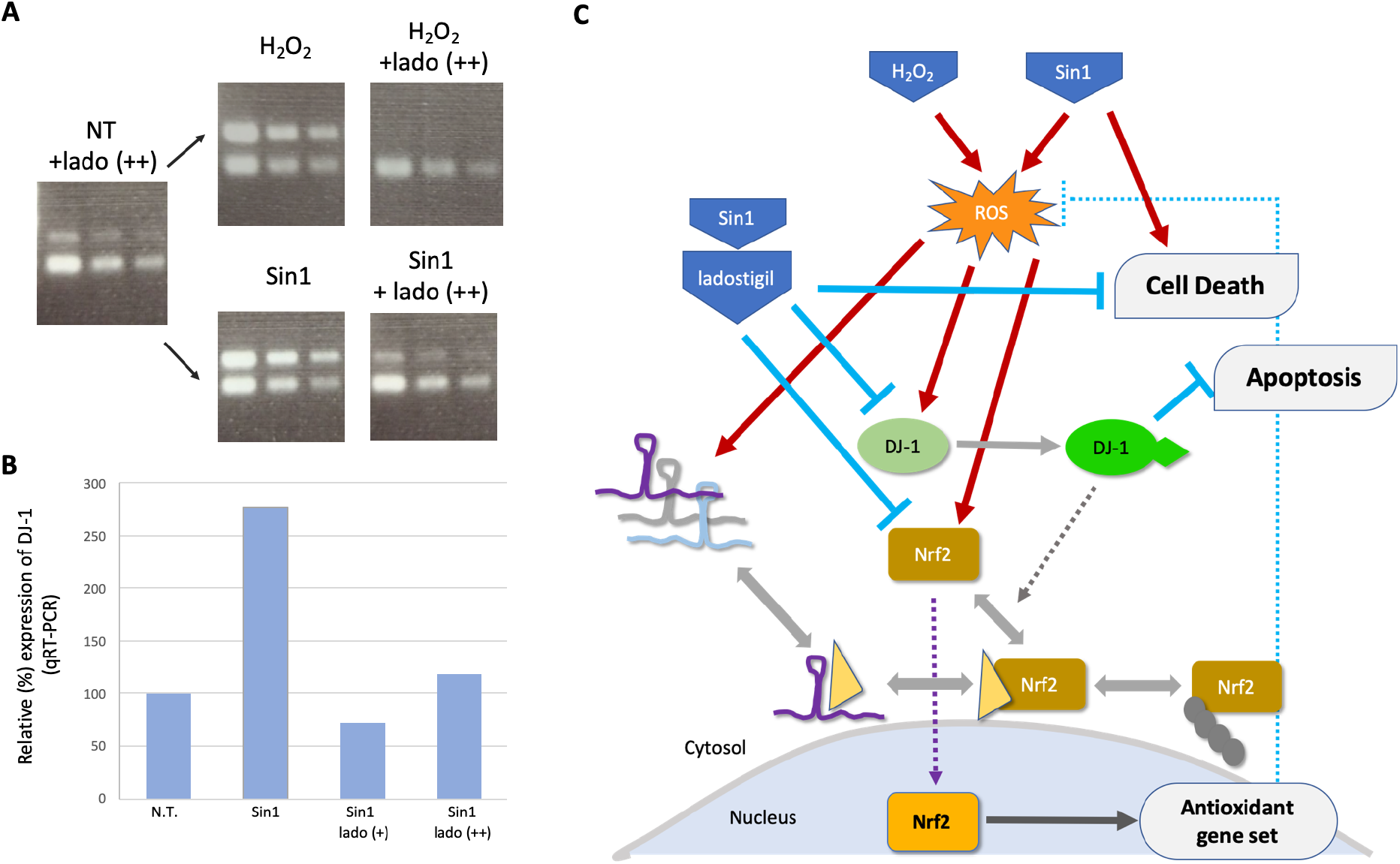
Expression levels of DJ-1 under oxidative stress conditions and ladostigil. **(A)** Semi-quantitative analysis of PCR amplicons of DJ-1 and β-actin. The amplicons were separated on 1.2% agarose gel. For each condition three lanes are shown representing loading of the PCR products that are equivalent to 1, 0.5 and 0.25 from left to right. cDNAs were prepared from cells at the indicated conditions for H_2_O_2_ and Sin1 treatment and following ladostigil at higher level (54 μM, marked as ++). **(B)** Results from a real-time quantitative PCR analysis of DJ-1 and β-actin of the untreated cells (NT), cells exposed to Sin1 (24 hours) with or without ladostigil at two concentrations. **(C)** Schematic diagram for major observations on oxidative stress regulation in SH-SY5Y cells. ROS production by H_2_O_2_ and Sin1 leads to enhanced cell death, upregulation of lncRNAs and induction of DJ-1 and Nrf2. Treatment with ladostigil, in combination of Sin1 results in the downregulation of DJ-1 and Nrf2, and inducescell death. Blue and red arrows indicate downregulation and upregulation, respectively. Dashed arrows indicate direct regulation. Nrf2 is depicted as bound to Keap1 (yellow triangle), ubiquitinated (gray circles) and translocated to nucleus.

The key findings in light of the oxidative stress generated by H_2_O_2_ and Sin1 (collectively labeled as ROS) and the effect of ladostigil in the presence of sustained Sin1 stimulation are summarized in **Figure 7C**. Sin1 upregulated a number of lncRNAs, of which the function of only a few were previously studied (e.g., MALAT1). DJ-1 serves as an oxygen sensor that shuttles between the cytosol, mitochondria, and the nucleus. Cytosolic DJ-1 restores Nrf2’s equilibrium and stabilizes it. By limiting the interaction with the suppressor Keap1, Nrf2 is stabilized. Specifically, by detaching it from the ubiquitination-induced protein complex that lead to its degradation. Unbound Nrf2 translocates to the nucleus and regulates the expression of a number of antioxidant enzymes through its binding to ARE (Antioxidant response elements) [55].

We show that ladostigil in the presence of Sin1 reduces DJ-1 and in Nrf2 expression levels relative to cells exposed to Sin1. In SH-SY5Y cells, ladostigil apparently reduces oxidative non-reversible damage, thus resulting in increased cell survival (**Figure 7C**). Note that a beneficial effect on apoptosis by increasing the activity of DJ-1 was reported in SH-SY5Y cells under an oxidation paradigm [56]. Finally, it was reported that Nrf2 is tightly regulated by a number of lncRNAs in the context of stress [57]. One such regulator of Nrf2 is MALAT1 which is a highly expressed ncRNA in SH-SY5Y cells and among the most significantly upregulated lncRNA induced by Sin1 (**Table 2**). The binding of MALAT1 to Keap1 regulates the antioxidant defense in diabetic retinopathy [58]. We highlight the lncRNAs as an understudied regulatory layer of cell physiology and stress-dependent response.

## 4. Discussion

In a previous study, the brain transcriptomes of young and old rats (6.5 and 22 months, respectively) were compared before and after chronic treatment with ladostigil. Performance of learning and memory tasks was improved in the ladostigil-treated rats compared to the control group. In the aging brain, RNA-seq analysis revealed a strong inflammatory signature associated with microglia [11]. The beneficial effect on the aged brain was attributed to ladostigil-induced suppression of the microglial activation [11,12]. In this work, we examined undifferentiated SH-SY5Y cells under oxidative stress conditions by inspecting their transcriptome. We focused primarily on the changes in the expression levels of lncRNAs [59-61]. Changes in the amounts and composition of ncRNAs (e.g., miRNAs, lncRNAs) are frequently observed in brain injury, ischemia, neuroinflammation, and pathophysiological processes [37,62].

Several unexpected discoveries emerged from the analysis of the lncRNA signature in SH-SY5Y under mild but continuous oxidative stress. Firstly, although ncRNAs accounted for 30% of all transcripts detected, only 6.7% (780) are expressed above a predetermined threshold (TMM >4, **Figure 3**). No function is associated with thousands of the ncRNA transcripts expressed at very low levels. Many lncRNAs were probably identified due to the high sensitivity of the RNA-seq technology [63] and do not necessarily reflect a robust transcriptional signal. Secondly, the majority of the Sin1-changed ncRNA transcripts are upregulated, with only a few that downregulated (**Figure 4**, Fisher exact test, p-value <1e0-5). Finally, the Sin1 upregulated lncRNAs’ absolute fold change (FC) is modest (FC, **Table 1**). At the molecular level, lncRNAs bind to (often unknown) targets which act on chromatin, sites of transcription, biological machines (e.g., spliceosome), competing for binding sites, recruiting or stabilizing proteins, and more. Despite a growing number of ncRNAs identified by advanced sequencing techniques [63], no function is associated with the vast majority of them. The function of only a handful of lncRNAs had previously been resolved (e.g., MALAT1, XIST).

Among the highly expressed lncRNAs (**Figure 3**), MALAT1, XIST, GAS5 (Growth arrest-specific 5), and NEAT1 (Nuclear paraspeckle assembly transcript 1) have been implicated in regulating epigenetic marks. Note that the genomic positions of lncRNAs are of a special importance. Many lncRNAs reside in chromosomal locations that overlap with miRNAs, suggesting that such lncRNAs may act as splicing regulators or antisense (i.e., competing with the gene where it resides in) [64]. For example, miR-3128 is found in the intron of NFE2L2 that encodes the Nrf2 transcription factor (TF). The lncRNAs are associated with expressed enhancers (eRNA), indirectly regulate the expression of neighboring genes. We identified a small number of ladostigil-upregulated genes that act as antisense or have eRNA-like properties (**Figure 6C, Table 2**). While there is no direct evidence for their function, specific lncRNA may have multiple functions (e.g., act both as eRNA and as ceRNA) through binding to different entities (e.g., DNA, protein) [65].

A direct role of miRNAs in maintaining SH-SY5Y cell homeostasis has been extensively studied [66]. In this study, we do not elaborate on the direct contributions of lncRNAs to post-transcriptional regulation by miRNAs. However, all SNHG genes are signified by their rich repertoire of miRNA binding sites (MBS). As a result, SNHGs may act by titrating miRNAs away from their targets, a mechanism known as ceRNA [67]. As a general theme, many lncRNAs act via sponging of specific miRNAs (e.g., [68,69]). For example, MALAT1 promotes oxygen-glucose deprivation/reoxygenation (OGD/R)-induced neuronal injury through ceRNA mechanisms [32,33]. Similarly, the beneficial effect of NORAD on cell survival was attributed to its miRNA sponging capacity [70].

The need to protect neurons from oxidative damage and proteotoxicity highlights the role of Nrf2 in the stress response (**Figure 7**). Nrf2 is a known master TF involved in redox homeostasis in response to oxidative stress. It is estimated that ∼3% of human coding genes are regulated by the Nrf2/Keap1 pathway [71]. In cells, Nrf2 fluctuates between alternative biochemical states that dictate the extent and kinetics of the stress response. We tested the impact of oxidative stressors on the expression level of DJ-1 (PARK7), and in the presence of ladostigil. DJ-1 is a multifunctional protein that connects various cell stressors and an upstream regulator of Nrf2 (**Figure 7**). It was shown that eliminating DJ-1 rendered SH-SY5Y susceptibility to cell death [72]. DJ-1 was confirmed to inhibit oxidative stress and consequently leads to cell protection [73,74]. As a fast-reacting oxidation sensor, the degree of its oxidized form (on Cys106) [75] is a proxy for a cell’s oxidative status. DJ-1 dominates the cellular fate of Nrf2 by impact on the stability of the Nrf2/Keap1 complex. Nrf2 strongly binds to its direct suppressor Keap1, makes it prone to polyubiquitin modification and degradation [76,77]. A crosstalk of DJ-1, with Nrf2 transcription, was demonstrated in protecting cells from oxidative stress by inducing the enzyme thioredoxin 1 (Trx1) by Nrf2 in in SH-SY5Y cells [78]. In accord with our findings, high expression of DJ-1 and Nrf2 occurs with cells exposed to oxidation insults. It was shown that overexpressed DJ-1 led to an increase in Nrf2 protein levels and the latter’s translocation to the nucleus. The enhanced binding of Nrf2 to ARE (e.g., at the Trx1 promoter) promotes transcription of a large spectrum of antioxidative enzymes (e.g., catalase, glutathione peroxidase). We postulate that the DJ-1/Nrf2 axis is as a hub for lncRNAs [79] and miRNAs [80] regulation. Notably, among miRNAs that directly target Nrf2 are several SH-SY5Y highly expressed ones (e.g., miR-93 and miR-27a) [57].

In this work, we studied undifferentiated SH-SY5Y cells. However, exposing naive cells to various combinations of growth factors and morphogenes (e.g., retinoic acid) drives differentiation, and the cells resemble mature dopaminergic-related neurons [23,81]. Differentiated SH-SY5Y cells become more vulnerable to oxidative damage [20]. In terms of human health, mutations in DJ-1 are linked to recessively inherited Parkinson’s disease. The mutated DJ-1 failed to act as an oxygen-based sensor. As a result, it disrupts DJ-1’s chaperone activity for preventing alpha-synuclein aggregation [82]. These DJ-1 multiple activities are not limited to brain injury, ischemia, or neurodegenerative diseases but also crucial in metabolic diseases (e.g., diabetes type 2 [83]). In a recent study, we showed that oxidative stress in SH-SY5Y undifferentiated cells leads to a coordinated alteration in the expression of genes that act in ER stress [29]. In the present study, we identified Sin1-induced lncRNAs that are known to regulate cell redox and ER homeostasis [51]. Based on our findings, we argue that lncRNAs act in concert to efficiently cope with parallel needs (e.g., damage to DNA, lipids, and proteins). MALAT1, for example, inhibited apoptosis via activating Nrf2 [35]. It also protects cells from oxidative stress by suppressing lipid peroxidation and damaged DNA [33] and directly binds Keap1, thus impacting Nrf2 TF availability.

In Sin1-stimulated cells, oxidative stress increased the expression of DJ-1 and Nrf2, whereas ladostigil prevented the induction of gene expression. We postulate that pre-incubation of ladostigil before exposure to oxidative stimuli protects cells from irreversible damage by lowering oxidative stress intensity [29]. Treatment with ladostigil leads to increased viability, which is likely mediated by a complex interplay involving lncRNAs, DJ-1, Nrf2 signaling, and the antioxidant response. This study suggests a framework in which lncRNAs coordinate between oxidative sensing mechanisms, the induction of antioxidant responses and cell decisions (e.g. apoptosis), where ladostigil partially diminishes these events by improving cell homeostasis.

## Abbreviations

AD: Alzheimer’s disease
CNS: central nerve system
DE: differentially expressed
ER: endoplasmic reticulum
FACS: fluorescence-activated cell sorting
FC: fold change
FCS: fetal calf serum
FDR: false discovery rate
H_2_O_2_: hydrogen peroxide
MBS: miRNA binding site
MEM: minimum essential medium
NT: not treated
Nrf2: nuclear factor erythroid 2-related factor 2
O_2_^−^: superoxide anion
ONOO^−^: peroxynitrite
PD: Parkinson’s disease
PI: propidium iodide
PS: phosphatidylserine
I/R: ischemia and reperfusion
RNS: reactive nitrogen species
ROS: reactive oxygen species
TF: transcription factor
TMM: trimmed mean of M-values

## Acknowledgments

We would like to deeply thank Prof. Marta Weinstock (The Hebrew University) for her constant support and supply of ladostigil. We thank Elyad Lezmi (The Hebrew University) for his contribution in the data processing and initial analysis. We thank Naomi van Wijk for reading the manuscript and the rest of the Linial’s lab members for useful comments and fruitful discussions.

